# RhoA Overexpression in Nucleus Accumbens-to-Ventral Tegmental Area Spiny Projection Neurons Induces Negative Affective-like Behavior

**DOI:** 10.64898/2026.11.22.768443

**Authors:** Payel Das, Samantha N. Scott, Daniela Franco, Cali A. Calarco, Ramesh Chandra, Joseph F. Cheer, Natalie E. Zlebnik, Shannon L. Cole, Mary Kay Lobo

## Abstract

**Rationale:** Major Depressive Disorder (MDD) is increasingly affecting populations worldwide and becoming a leading cause of psychological disabilities. However, MDD patients do not respond to current medications, resulting in treatment resistance. This situation demands more basic research to identify targetable molecules and the specific brain circuits and cell types that underlie negative affective pathology. Ras homolog family member A (RhoA) mediates negative affective-like behaviors via dendritic atrophy of nucleus accumbens (NAc) D1 (dopamine receptor 1)-Spiny Projection Neurons (SPNs), making RhoA a promising target for new depression therapeutics. However, the sufficiency of RhoA elevation in NAc projection-specific D1-SPNs is yet to be investigated.

**Objectives:** Whether RhoA overexpression selectively in NAc-to-ventral tegmental area (VTA) SPNs (NAc-VTA SPNs) is sufficient to induce negative affective-like behavior. Whether chronic stress (CS) induces RhoA mRNA expression in this specific projection.

**Methods:** A retrograde (Rg) vector expressing Cre recombinase was infused into the VTA together with Cre-dependent vectors overexpressing either a control expressing only enhanced Yellow Fluorescent Protein (eYFP) or RhoA-eYFP in NAc-VTA neurons. Mice then underwent a battery of affective assays (social interaction, etc.) and a two-choice sucrose operant task assessing motivation. RNA in situ hybridization (RHA-ish) quantified RhoA mRNA in NAc-VTA D1-SPNs of chronically stressed versus unstressed (control) mice.

**Results:** RhoA overexpression in NAc-VTA SPNs, which are primarily D1-SPNs, elicited negative affective-like phenotypes, including social avoidance, reduced open-arm exploration, decreased grooming, and increased latency to approach food. RhoA and control groups acquired a food operant task similarly. However, RhoA overexpression significantly reduced operant responding and breakpoint during progressive-ratio testing. RhoA overexpression in the NAc-VTA projection neurons also diminished persistent sucrose-seeking behavior. Chronic stress induced RhoA mRNA expression selectively in NAc-VTA neurons.

**Conclusion:** These studies identify a potential NAc D1-SPN circuit that underlies negative affective behaviors through RhoA mechanisms.

## Introduction

Major depressive disorder (MDD) is the predominating cause of psychological disability worldwide (Murray and Lopez 1997; Zhang et al. 2026; Du et al. 2026; Qin and Wei 2026); more than two-thirds of individuals continue to exhibit lasting symptoms despite treatment, and up to 40% exhibit resistance to conventional therapies (Al-harbi 2012; Dodd et al. 2021; Touloumis 2021; Havlik et al. 2024; Siyoum et al. 2024; Fiorillo et al. 2025). Chronic stress is a major risk factor for MDD (Turner and Beiser 1990; McGonagle and Kessler 1990; Hammen et al. 2009; Wang et al. 2024; Malykhin et al. 2025), which is marked by impaired motivation (Treadway and Zald 2011), reduced pursuit of rewards (Daniels et al. 2025), and impaired decision-making (Treadway et al. 2012), contributing to functional disability (Cléry-Melin et al. 2019; Halahakoon et al. 2020) and worse quality of life (Wong et al. 2024). Motivation, which guides organisms toward resources (e.g., food) and away from threats (Brown 1948; Lang and Bradley 2010; Bodas et al. 2026; Flores et al. 2026), is regulated by reward circuitry (Lobo and Nestler 2011; Gutierrez et al. 2011; LeGates et al. 2018; Illescas-Huerta et al. 2026). Chronic stress can disrupt reward circuits, contributing to depression-related phenotypes (McEwen 2007, 2008, 2012, 2017; McEwen and Gianaros 2010; Russo and Nestler 2013); understanding how it does so (Fox and Lobo 2019) may identify therapeutic targets.

In murine models, chronic social defeat stress (CSDS) (Krishnan et al. 2007; Golden et al. 2011) and chronic witness defeat stress (CWDS) (Warren et al. 2013; Sial et al. 2016; Iñiguez et al. 2018) produce negative affective-like behaviors resembling aspects of depression: stress-susceptible animals show social avoidance (low-interactors) and reduced sucrose preference compared to stress-resilient (high-interactors) mice (Krishnan et al. 2007; Golden et al. 2011; Warren et al. 2013; Sial et al. 2016; Iñiguez et al. 2018). These models allow investigation of stress-induced affective and motivational alterations (Deussing 2006; Krishnan and Nestler 2011; Von Muecke-Heim et al. 2021; Von Mücke-Heim et al. 2023). Distinct neuronal populations within the nucleus accumbens (NAc) have been implicated (Russo and Nestler 2013; Fox and Lobo 2019). Decoding how stress alters their molecular and cellular processes may clarify stress-induced changes in motivated behavior.

In the mesocorticolimbic reward system, the NAc is an integral hub in appetitive motivation (Carlezon and Thomas 2009; Volman et al. 2013), including the pursuit of natural (Berridge and Kringelbach 2015) and drug (Nestler and Lüscher 2019) rewards. Chronic stress induces persistent neuroplasticity in the NAc (Golden et al. 2013; Francis et al. 2015; Hamel et al. 2017; Fox et al. 2020b; Soares-Cunha et al. 2020; Franco et al. 2022; Zhou et al. 2022), implicating it in anhedonia (Lim et al. 2012; Daniels et al. 2025) and social avoidance (Francis and Lobo 2017). Spiny projection neurons (SPNs) expressing dopamine receptor 1 (D1) or 2 (D2) (Lobo and Nestler 2011; Kupchik et al. 2015) make distinct contributions to reward-and stress-related behaviors (Smith et al. 2013; Francis et al. 2015; Kai et al. 2015; Soares-Cunha et al. 2020; Pagliusi et al. 2022; Correia et al. 2023; Faust et al. 2025; Carter et al. 2025; Siemonsmeier et al. 2026). NAc D1-SPNs are associated with stress-induced behavioral adaptations and exhibit robust synaptic plasticity following chronic stress (Lim et al. 2012; Lobo et al. 2013; Francis et al. 2015, 2017, 2019; Francis and Lobo 2017; Chandra et al. 2017; Fox et al. 2020a; Kumar et al. 2026; Burek and Carlezon 2026).

Our laboratory identified RhoA signaling in NAc D1-SPNs as a cell-type- and region-specific substrate of chronic stress ( Francis et al. 2019; Fox et al. 2020a). CSDS produces dendritic atrophy in NAc D1-SPNs, but not D2-SPNs (Fox et al. 2020b), which correlates with social avoidance (Fox et al. 2020a) and accompanies increased RhoA and its downstream effector Rho-kinase (ROCK) signaling (Fox et al. 2020a). Manipulating RhoA-ROCK signaling alters D1-SPN structure and behavior: RhoA activation reduces dendritic complexity and produces negative affective-like behaviors without stress, whereas RhoA/ROCK inhibition prevents stress-induced behavioral changes (Francis et al. 2019; Fox et al. 2020a). RhoA also regulates D1-SPN functional adaptations to stress (Fox and Lobo 2019; Fox et al. 2020a), including excitability and spine density (Francis et al. 2019). Because NAc D1-SPNs comprise multiple projection-defined populations (Kupchik et al. 2015; Baimel et al. 2019; Pardo-Garcia et al. 2019), it remains unknown whether projection-specific RhoA overexpression (OE) induces negative affective-like behaviors. The NAc-VTA pathway, through which D1-SPNs provide GABAergic input to VTA neurons (Xia et al. 2011; Bocklisch et al. 2013; Edwards et al. 2017; Simmons et al. 2017; Yang 2018; Polter et al. 2018; Khayat and Yaka 2024), may shape stress-related social interaction, reward sensitivity, and anhedonia (Qi et al. 2022). Manipulating NAc D1-SPNs alters social avoidance and sucrose preference in chronic stress as well has other negative affective or exploratory behaviors in stress naïve subjects (Francis et al. 2015, 2017, 2019; Francis and Lobo 2017; Fox et al. 2020a). However, whether NAc-VTA projection -specific neurons contribute to theses behavioral outcomes remains unknown. To address this, the present study investigated RhoA signaling in NAc-VTA SPNs as a circuit-specific mechanism of negative affect, exploratory and effort-based motivation behavioral outcomes.

## Methods

### Animals and experimental design

All procedures were approved by the Institutional Animal Care and Use Committee of the University of Maryland School of Medicine and followed federal guidelines for the humane care and use of laboratory animals. Male and female C57BL/6J mice (The Jackson Laboratory) were housed 3-5 per cage on a 12-h light/dark cycle with food and water available ad libitum. We used 36 (18 male, 18 female) for the behavioral battery, 33 (16 male, 17 female) for sucrose operant testing, and 48 mice (25 male, 23 female) for RNAscope.

### Stereotaxic surgery

Mice were anesthetized with isoflurane (4% induction, 1-2% maintenance) and secured in a stereotaxic frame. Using 33-gauge Hamilton syringes, mice received bilateral infusions of AAV-Rg-Cre into the VTA (AP -3.2, ML +/-1.0, DV -4.6 mm, 7-degree angle) and AAV2.2-Ef1a-DIO-RhoA-eYFP or Ef1a-DIO-eYFP into the NAc (AP +1.6, ML +/-1.5, DV -4.4 mm, 10-degree angle), 0.4 microliters per side. Plasmid for AAV2.2-Ef1a-DIO-RhoA-eYFP was constructed from pcDNA3RhoA plasmid (Addgene #12965), inserted into an Ef1a-DIO-eYFP vector following established protocols (Fox et al. 2020a). For the behavioral battery, 19 mice (9 male, 10 female) received RhoA-OE, and 17 (9 male, 8 female) received eYFP; for the operant task, 17 (8 male, 9 female) received RhoA-OE, and 16 (8 male, 8 female) received eYFP. Mice recovered for three weeks before testing to allow viral expression.

### Chronic social stress paradigms

CSDS and CWDS were conducted as previously described, with adaptations for vicarious resident-intruder stress. Male C57BL/6J intruders were exposed to a dominant CD-1 resident for 10 min daily in a hamster cage divided by a perforated partition. Only males underwent CSDS, because CD-1 residents do not reliably attack females without further modification. For CWDS, a male or female mouse observed the same encounter from the opposite side of the partition for 10 min. After each session, the defeated mouse remained housed across the divider from its resident, and the witness was relocated to observe a new pairing, giving 10 unique CD-1 pairings across 10 consecutive days. Control mice were pair-housed across a perforated divider with a sex-matched conspecific for the same period. All mice were singly housed immediately after the final session. (Golden et al. 2011; Sial et al. 2016; Iñiguez et al. 2018)

### Three-chamber social interaction test (3CSI)

Twenty-four hours after the final stress session, mice were tested in a three-chamber arena (60 x 40 cm) divided into three equal compartments by perforated clear acrylic partitions. Mice explored the arena with two empty wire mesh cups for 5 mins. For next 5 mins’ exploration, a novel sex-matched conspecific was placed in one cup, placed in the interaction zone while the other cup in the isolation zone left empty. Mice spending less than the threshold time in the social chamber were classified as Low Social Interactors (LOW) and those spending more as High Social Interactors (HIGH). The threshold is corresponded to the average time that unstressed control (sex-matched) animals spent in the interaction zone. (Franco et al. 2022)

### Open field test (OFT)

Mice were placed in a 43 x 43 x 43 cm non-porous arena and recorded for a single session of 15 mins. Total distance traveled indexed locomotor activity, and time spent in the central versus outer zone indexed anxiety-like avoidance. (Seibenhener and Wooten 2015; Kraeuter et al. 2019)

### Sucrose splash test (SST)

Mice were placed individually in a 4000 mL glass beaker, and the dorsal fur was sprayed three times with 10% sucrose solution (w/v). Grooming was recorded for 5 min with ANY-maze and quantified from video by a blinded experimenter using Kinescope 3.0.4. (Kokras et al. 2017; Strekalova et al. 2023)

### Elevated zero maze test (EZM)

The maze (San Diego Instruments, San Diego, California, US) is a black acrylic annulus 105 cm in diameter with a 10 cm wide track elevated 72 cm above the floor, divided into two open quadrants bordered by 1 cm lips and two closed quadrants enclosed by 28 cm walls. Each mouse was placed in a closed quadrant and recorded for 5 min under constant lighting. The maze was cleaned with 70% ethanol between trials. (Shepherd et al. 1994)

### Novelty-suppressed feeding test (NSFT)

Mice were food-restricted overnight and excluded if they lost more than 20% of body weight. Testing was conducted in a 60 x 40 cm arena lined with approximately 2 cm of woodchip bedding, with a single food pellet placed on white paper at the center. Each mouse was introduced at the same corner, and latency to the first bite was recorded over 15 min; mice that did not eat were assigned the maximum latency of 900 s. Mice were then transferred to a test cage containing a fresh pellet, and latency to contact the food was recorded for up to 5 min. (Santarelli et al. 2003; Samuels and Hen 2011)

### Sucrose operant behavior

Testing was conducted in operant chambers (Med Associates, VT, USA) fitted with two nose-poke ports, a cue light, and a house light. Mice were trained on a fixed ratio 1 (FR1) schedule for four 90-min sessions (one per day). An active nose poke delivered a 14 mg sucrose pellet (Bio-Serv, NJ, USA) and illuminated the cue light above the active port, followed by a 10-s timeout during which pokes produced no reward; the house light then signaled reward availability. Inactive pokes produced no cue or pellet, and active and inactive ports were counterbalanced across mice. Mice then completed three FR3 sessions. Next, incentive motivation was assessed under a progressive ratio (PR) schedule in which the requirement increased as responses per reinforcer = 5e^(0.2 x reinforcer number) -5 (Richardson and Roberts 1996), with sessions ending after 20 min without a response and cues presented as in FR testing; mice were returned to FR3 for two sessions afterwards. Twenty-four hours after the final FR3 session, mice underwent a 90-min cue-induced sucrose-seeking test in which active pokes produced the sucrose-paired cue but no pellet. Mice then remained in the vivarium for 30 days before an identical second sucrose-seeking test.

### RNA in situ hybridization

Brains were rapidly extracted, flash-frozen in isopentane at -40 degrees C for 20 s, and stored at -80 degrees C. Coronal sections (16 micrometers) containing the NAc were cut on a cryostat (Leica, Wetzlar, Hesse, Germany), mounted on Superfrost Plus slides, and stored at -80 degrees C. RNAscope was performed with the Multiplex Fluorescence Reagent Kit v1 (Advanced Cell Diagnostics, Newark, California, USA) according to the fresh-frozen protocol and previously published methods (Wang et al. 2012). Sections were fixed in 10% neutral buffered formalin for 20 min, treated with protease IV for 20 min, and hybridized with probes against Drd1 (custom channel 3, Catalog no. 519411-C3), Cre (Catalog no. 461901), and RhoA (Catalog no. 409431-C2). Signal was developed with Amp 4 Alt B-FL, and images were acquired at 63x on a Leica SP8 confocal microscope (Leica, Wetzlar, Hesse, Germany). RhoA expression was quantified in QuPath v0.5.1; (Bankhead et al. 2017) by an experimenter blinded to experimental groups.

### RT-qPCR

Bilateral NAc (four 14-gauge punches per mouse) and VTA (one 12-gauge punch per mouse) punches were collected 24 h after the final social interaction and stored at -80⁰C. Total RNA was isolated with TRIzol (Invitrogen, 15596018), purified with the EZNA MicroElute Total RNA kit (Omega Bio-Tek, R6831-01), including DNase treatment (Catalog no. 79254, QIAGEN N.V., Venlo, Limburg, Netherlands) and assessed on a NanoDrop 1000 (Thermo Fisher Scientific, Waltham, Massachusetts, USA). cDNA was synthesized from 400 ng RNA (iScript, Bio-Rad 1708891) and diluted to 2 ng/uL. RT-qPCR was performed using PerfeCTa SYBR Green FastMix (Quantabio, Catalog no. 95072, Beverly, Massachusetts) on a Bio-Rad CFX384 system (Bio-Rad Laboratories Inc., Hercules, California, USA) (Calarco et al. 2021). Relative expression was calculated by the 2^-DDCt method (Livak and Schmittgen 2001) with GAPDH as reference and normalized to sex-matched unstressed controls. Primers: RhoA forward 5’-TCTTCAGCAAGGACCAGTTC-3’, RhoA reverse 5’-TCCCACAAAGCCAACTCTAC-3’; eYFP forward 5’-AGCAGAAGAACGGCATCAA-3’, eYFP reverse 5’-GGGTGTTCTGCTGGTAGT-3’.

### Statistics

Statistical analyses were performed using GraphPad Prism 11.0.0 (www.graphpad.com) or IBM SPSS Statistics 31.0.0.0 (www.ibm.com/products/spss-statistics), as indicated. For social interaction, open field, elevated zero maze, sucrose splash, novelty feeding tests, and qPCR mRNA analyses (DCt values), data were analyzed using ordinary two-way ANOVA with sex (male vs. female) and AAV treatment (eYFP vs. RhoA-OE) as between-subjects factors. Significant main effects or interactions were assessed using Šidák’s multiple-comparisons test or Benjamini-Krieger-Yekutieli test (BKY, as recommended by the software), with a single pooled variance, in GraphPad Prism 11.0.0. *Sexes are combined into each of treatment group where sex-effects are not seen*. Sucrose self-administration data were analyzed using IBM SPSS Statistics 31.0.0.0. For FR1, FR3, and PR schedules, active and inactive nose poke responses and sucrose rewards earned were analyzed using mixed-design ANOVAS, with session day as a within-subjects factor and AAV treatment and sex as between-subjects factors. For FR1 acquisition, active and inactive nose poke responses were analyzed with port (active vs inactive) included as an additional within-subjects factor. For PR responding, breakpoint was analyzed across sessions using a mixed-design ANOVA with session day as within-subjects factor and AAV treatment and sex as between-subjects factors. Cue-induced sucrose-seeking responses following abstinence were analyzed using a two-way ANOVA with AAV treatment and sex as a between-subjects factors, and abstinence duration (24-hour vs 30-day) as a within-subjects factor. Significant main effects and interactions were followed by Bonferroni-adjusted pairwise comparisons or Tukey’s multiple-comparisons test, as appropriate for the analysis. For the RNAscope-in situ hybridization (*ISH)* experiments, one-way ANOVA was used to compare control, Low Social Interactor, and High Social Interactor groups, followed by Tukey’s multiple-comparisons test in GraphPad Prism 11.0.0. For the correlational study between RhoA-mRNA level vs. the time in the interaction zone, simple linear regression was performed. Potential outliers were identified using Grubbs’ test (Grubbs 1950, 1969) in GraphPad Prism and were excluded from subsequent analyses when statistically identified as outliers. Statistical significance was defined as p < 0.05. Data are presented as mean ± SEM.

### Artificial Intelligence (AI)

Generative AI was not used to create the content of this manuscript. However, Grammarly was used for grammar corrections in the initial draft by the first author. Consensus was used for citation identification but manually verified. Claude was used for constructive feedback by simulating an expert reviewer with a prompt generated by the author.

## Results

### RhoA overexpression in NAc-VTA D1-SPNs was confirmed by RT-qPCR, and Cre expression was enriched for D1-SPNs

To selectively manipulate RhoA expression in NAc-VTA SPNs, Cre-dependent viral constructs – either eYFP (for control group) or RhoA-eYFP (for RhoA-OE group) was injected into the NAc in combination with retrograde Cre into the VTA (**Fig. 1A**). Representative fluorescence images confirmed expression of DIO-RhoA-eYFP/ DIO-eYFP in the NAc successful AAV-targeting and circuit-specific RhoA-OE (**Fig. 1B**). RT-qPCR analysis of NAc tissue punches confirmed robust RhoA-OE in mice receiving AAV-DIO-RhoA-eYFP compared to eYFP controls (**Fig. 1C**; treatment: F(1,33) = 192.7, p < 0.0001), with Šidák post-hoc comparisons confirming significantly higher expression in RhoA-OE mice compared to eYFP controls (p < 0.001). There was no main effect of sex (F (1, 33) = 0.04, p=0.8500) and no sex by treatment interaction (F (1, 33) = 0.04, p=0.8501). Confocal imaging after multiplex RNA-ish analysis confirmed the cell-type specificity of the target pathway: expressing both Drd1 and Cre (D1+/Cre+, marked by yellow arrowhead). (**Fig. 1D**). Quantification of Drd1 and Cre mRNA transcripts revealed that approximately 99.2% (Mean ± SEM) of VTA-projecting NAc neurons were D1-SPNs, demonstrating a high degree of colocalization

**Figure 1.**
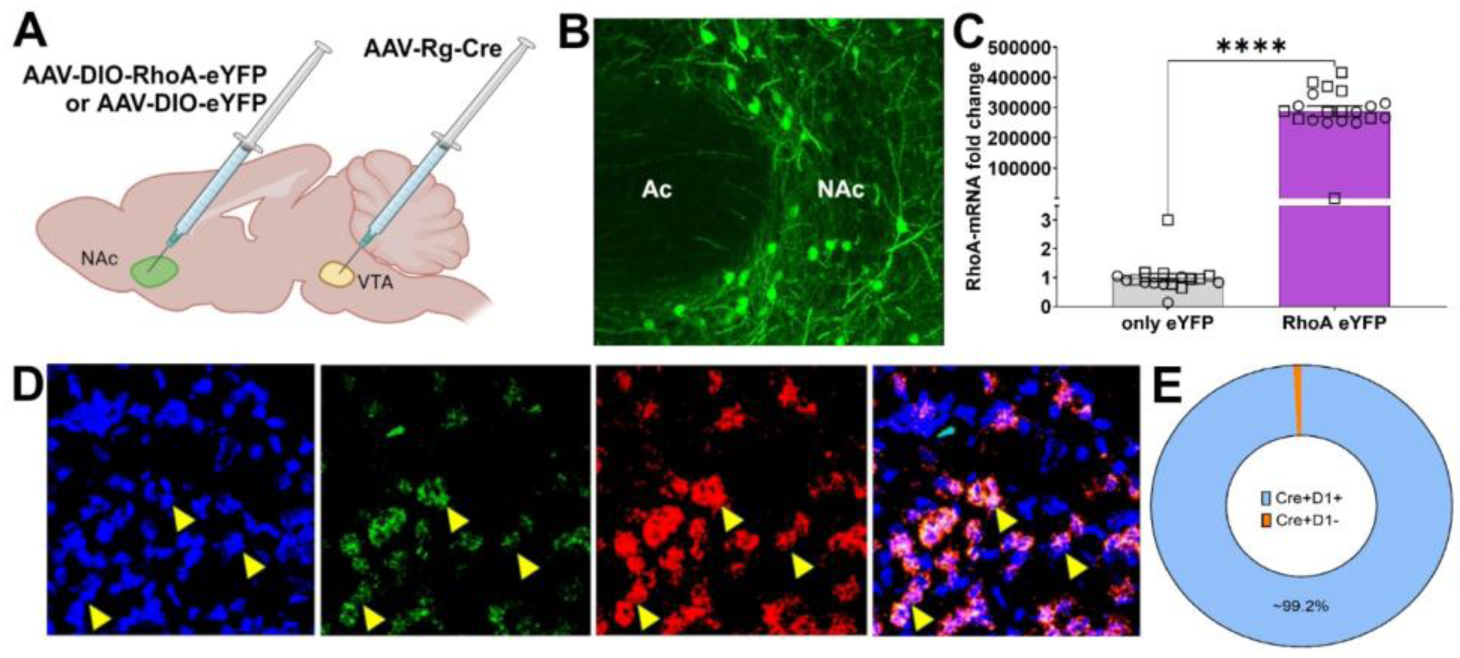
RT-qPCR confirmed RhoA overexpression in NAc and RNAscope showed Cre expression was selective to D1-SPNs. (A) Viral strategy uses Cre-lox system to express RhoA-eYFP or eYFP strictly in NAc-VTA neurons by injecting AAV-Rg-Cre in VTA (Rg: retrograde) and AAV-DIO-RhoA-eYFP or AAV-DIO-eYFP in NAc. (B) Representative image, followed by immunohischemistry against eYFP on NAc slices showed eYFP expression in the NAc, Ac: anterior commissure. (C) RT-qPCR against NAc tissue punches showed increased RhoA-mRNA expression (fold change) in the mice that received AAV-DIO-RhoA-eYFP compared to those that received AAV-DIO-eYFP. (D) Representative image, followed by RNA-ish on NAc slices showed DAPI (nucleus, blue), Cre (green), and Drd1 (D1, red) in the NAc, along with the merged image showing yellow arrowheads marked cells as Cre⁺/Drd1⁺; light-blue arrowheads marked cells as Cre⁻/Drd1⁺. (E) Approximately 99.2% of NAc-VTA neurons (Cre-positive) are Drd1-positive. (Squares: males, circles: females; all figures (where applies) represent mean ± SEM, statistical significance was assessed using p = 0.05, using p = 0.05, *p < 0.05, **p < 0.01, ***p < 0.001, and ****p < 0.0001; ns, not significant.) between Cre-positive and Drd1-positive puncta, with minimal overlapping Cre-expression in Drd1-negative cells (Fig. 1E).

### RhoA-OE in NAc-VTA SPNs induces negative affective-like behavior in mice, in absence of chronic **social stress.**

To determine if RhoA overexpression (RhoA-OE) in NAc-VTA SPNs induces negative affective-like behaviors and disrupts exploratory behavior, we performed a battery of behavioral tests in mice expressing RhoA in this projection-defined population. In brief, AAV-retrograde Cre was injected into the VTA, and Cre-dependent AAV-DIO-RhoA-eYFP or AAV-DIO-eYFP, used as a vector control, was injected in the NAc of wildtype mice (**Fig. 2A**). Following a 3-week period for viral expression, mice underwent a battery of behavioral tests, in specific order (**Fig. 2A**).

**Figure 2.**
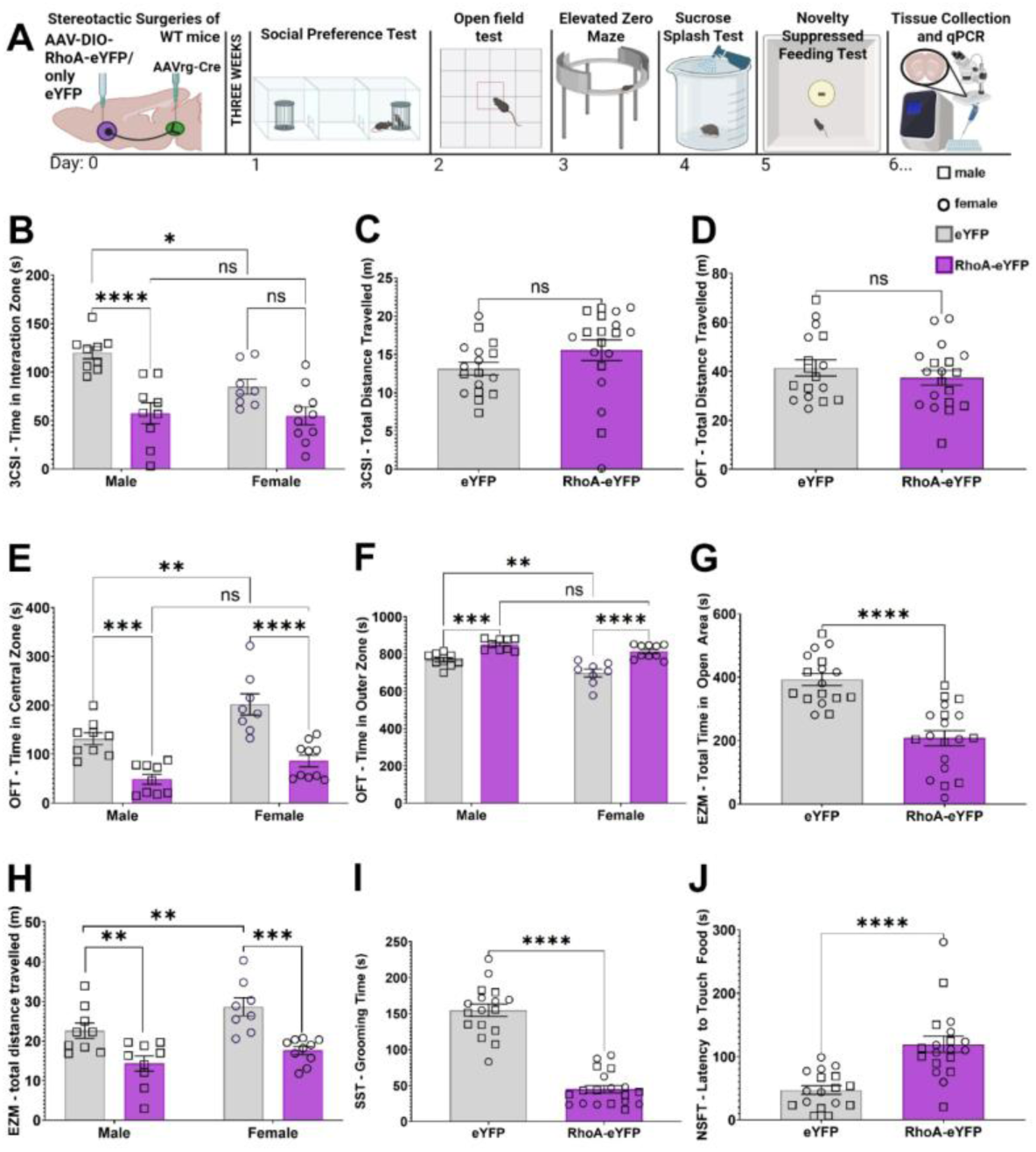
RhoA-OE in NAc-VTA D1-SPNs induced negative affective-like behaviors in the absence of chronic social stress. (A) Experimental timeline. AAV-Rg-Cre was injected into the VTA, and AAV-DIO-RhoA-eYFP or AAV-DIO-eYFP was injected into the NAc of wild-type mice. Following a 3-week viral-expression (recovery) period, mice underwent the three-chamber social interaction test (3CSI; day 1), open field test (OFT; day 2), elevated zero maze (EZM; day 3), sucrose splash test (SST; day 4), and novelty suppressed feeding test (NSFT; day 5), followed by tissue collection for RT-qPCR (day 6). (B) RhoA-OE mice spent less time in the interaction zone of the 3CSI than eYFP controls (post-hoc difference significant in males, not in females), and eYFP males spent more time in the interaction zone than eYFP females. (C) Total distance traveled during the habituation phase of the 3CSI did not differ between groups. (D) Total distance traveled in the OFT did not differ between groups. (E, F) RhoA-OE mice spent less time in the central zone (E) and more time in the outer zone (F) of the OFT arena than eYFP controls; eYFP females spent more time in the central zone and less time in the outer zone than eYFP males. (G) RhoA-OE mice spent less time in the open area of the EZM than eYFP controls. (H) RhoA-OE mice traveled less distance in the EZM than eYFP controls, and eYFP females traveled more than eYFP males. (I) RhoA-OE mice spent less time grooming in the SST than eYFP controls. (J) RhoA-OE mice showed a longer latency to touch the food in the NSFT than eYFP controls. (Squares: males, circles: females; gray bars: eYFP, purple bars: RhoA-eYFP; all figures represent mean ± SEM, statistical significance was assessed using p = 0.05, *p < 0.05, **p < 0.01, ***p < 0.001, and ****p < 0.0001; ns, not significant.)

In the three-chamber social interaction test, the main effect of treatment showed that RhoA-OE mice spent significantly less time in the interaction zone than eYFP controls, indicating reduced social interaction (**Fig. 2B**; F (1, 32) = 27.89, p<0.0001; Šidák, Male-eYFP vs. Male-RhoA-eYFP: p<0.0001). A significant main effect of sex was also observed (**Fig. 2B**; F (1, 32) = 4.65, p=0.0387, Šidák, Male-eYFP vs. Female-eYFP: p=0.0377), with males spending significantly more time in the interactions zone than females. The sex by treatment interaction did not reach significance (F (1, 32) = 3.42, p=0.0736).

In contrast, neither locomotor activity during the habituation phase of the three chamber social interaction test (**Fig. 2C**; treatment: F (1, 32) = 2.17, p=0.1510; sex: F(1,32) = 0.03, p = 0.85; interaction: F(1,32) = 1.32, p = 0.2586) nor total locomotor activity in the open-field test (**Fig. 2D**; treatment: F(1,32) = 1.00, p = 0.3239; sex: F(1,32) = 1.86, p = 0.1824; interaction: F(1,32) = 1.52, p = 0.2264) was affected by RhoA-OE, suggesting that the reduction in social interaction was not attributable to altered locomotor activity.

In the open-field test, a main effect of treatment indicated that RhoA-OE mice spent significantly less time in the center zone (**Fig. 2E**; F (1, 32) = 50.08, p<0.0001; Šidák, Male-eYFP vs. Male-RhoA-eYFP: p = 0.0008; Šidák, female-eYFP vs. female-RhoA-eYFP: p<0.0001) and more time in the outer zone (**Fig. 2F**; F (1, 32) = 50.08, p<0.0001; Šidák; Male-eYFP vs. Male-RhoA-eYFP: p = 0.0008; female-eYFP vs. female-RhoA-eYFP: p <0.0001) than eYFP controls. A significant main effect of sex for both center and outer zone (**Fig. 2F, G**; F (1,32) = 14.72, 14.72 p = 0.0006, 0.0006, respectively) and outer zone, revealed that females spent more time in the center zone and less time in the outer zone than the males (**Fig. 2E**, **2F**; F (1, 32) = 14.72, 14.72 p=0.0006, 0.0006; respectively; Šidák for both, Male-eYFP vs. Female-eYFP: p = 0.0065). There was no significant sex by treatment interaction (F (1,32) = 1.36, p = 0.2524).

A similar behavioral pattern was observed in the elevated zero maze (EZM). We observed a significant main effect of treatment on time spent in the open area (**Fig. 2G**; F (1, 32) = 34.18, p<0.0001; Šidák, Male-eYFP vs. Male-RhoA-eYFP: p = 0.0089; female-eYFP vs. female-RhoA-eYFP: p <0.0001), with RhoA-OE mice spending significantly less time in the open areas than eYFP controls (Fig. 2I, J; F(1,32) = 34.18, 23.01, p < 0.0001, respectively). There were no significant main effects of sex (open: F (1, 32) = 0.0036, p=0.9523; closed: F (1, 32) = 0.13, p=0.7185) and no significant sex by treatment interactions (open: F (1, 32) = 1.35, p=0.2541; closed: F (1, 32) = 1.70, p=0.2014).

For total distance traveled in the EZM, the analysis revealed significant main effects of treatment (F (1, 32) = 29.18, p<0.0001; BKY, Male-eYFP vs. Male-RhoA-eYFP: p=0.0013; Female-eYFP vs. Female-RhoA-eYFP: p=0.0001), reflecting a reduction in locomotor activity in RhoA-OE mice compared to eYFP controls (**Fig. 2H**). There was also a significant main effect of sex (F (1, 32) = 6.67, p=0.0146; p=0.0099; BKY, Male-eYFP vs. Female-eYFP: p=0.0099) driven by female mice traveling more than male mice only in the control group (**Fig. 2H**). No significant sex by treatment interaction was observed for total distance (F (1, 32) = 0.57, p=0.4552). The reduction in EZM locomotor activity may reflect increased occupancy of the closed areas, although a contribution of reduced locomotion cannot be excluded. Together, these findings indicate increased avoidance of exposed areas following RhoA-OE, consistent with an anxiety-like phenotype.

In the sucrose splash test, a significant treatment effect on grooming behavior, with RhoA-OE mice exhibiting significantly less grooming duration than eYFP controls (**Fig. 2I**; F(1,32) = 128.3, p < 0.0001, Šidák, Male-eYFP vs. Male-RhoA-eYFP: p<0.0001; Female-eYFP vs. Female-RhoA-eYFP: p<0.0001). This finding may reflect a reduction in self-care motivational drive or an increase in negative affective-like behavior following RhoA-OE. There was no significant main effect of sex (F(1,32) = 1.89, p = 0.1783) and no significant sex by interaction (F(1,32) = 1.29, p = 0.2647) observed for grooming behavior.

Finally, in the novelty suppressed feeding test, RhoA-OE mice exhibited significantly greater latency to approach and contact the food compared with control mice, as evidenced a significant main effect of treatment (**Fig. 2J**; F(1,31) = 21.18, p < 0.0001, Šidák, Male-eYFP vs. Male-RhoA-eYFP: p=0.0068; Female-eYFP vs. Female-RhoA-eYFP: p=0.0171), consistent with increased avoidance and enhanced anxiety-like behavior in a novel environment. There was no effect of sex (F(1,31) = 2.12, p = 0.1557) nor sex by interaction effect (F(1,31) = 0.033, p = 0.8578).

Collectively, RhoA-OE in NAc-VTA SPNs produced a behavioral profile characterized by reduced social interaction, increased avoidance or exposed areas, reduced grooming, and increased latency to approach food in a novel environment. These effects occurred in the absence of generalized locomotor impairment in the three-chamber habituations and open-field tests, although locomotor activity was reduced in the EZM. Together these findings indicate that RhoA-OE in NAc-VTA SPNs is sufficient to induce negative affective-like and anxiety-related behavioral phenotypes in the absence of chronic social stress.

### NAc-VTA D1-SPN RhoA overexpression blunts sucrose self-administration

Following three weeks of viral incubation, mice underwent sucrose self-administration training followed by sucrose-seeking tests after a period of acute or chronic abstinence (**Fig. 3A**). All behavioral data was analyzed using a three-way mixed-model ANOVA, with AAV treatment and sex as between-subjects factors and port or training day as within-subjects factors) selected according to the behavioral test.

**Figure 3.**
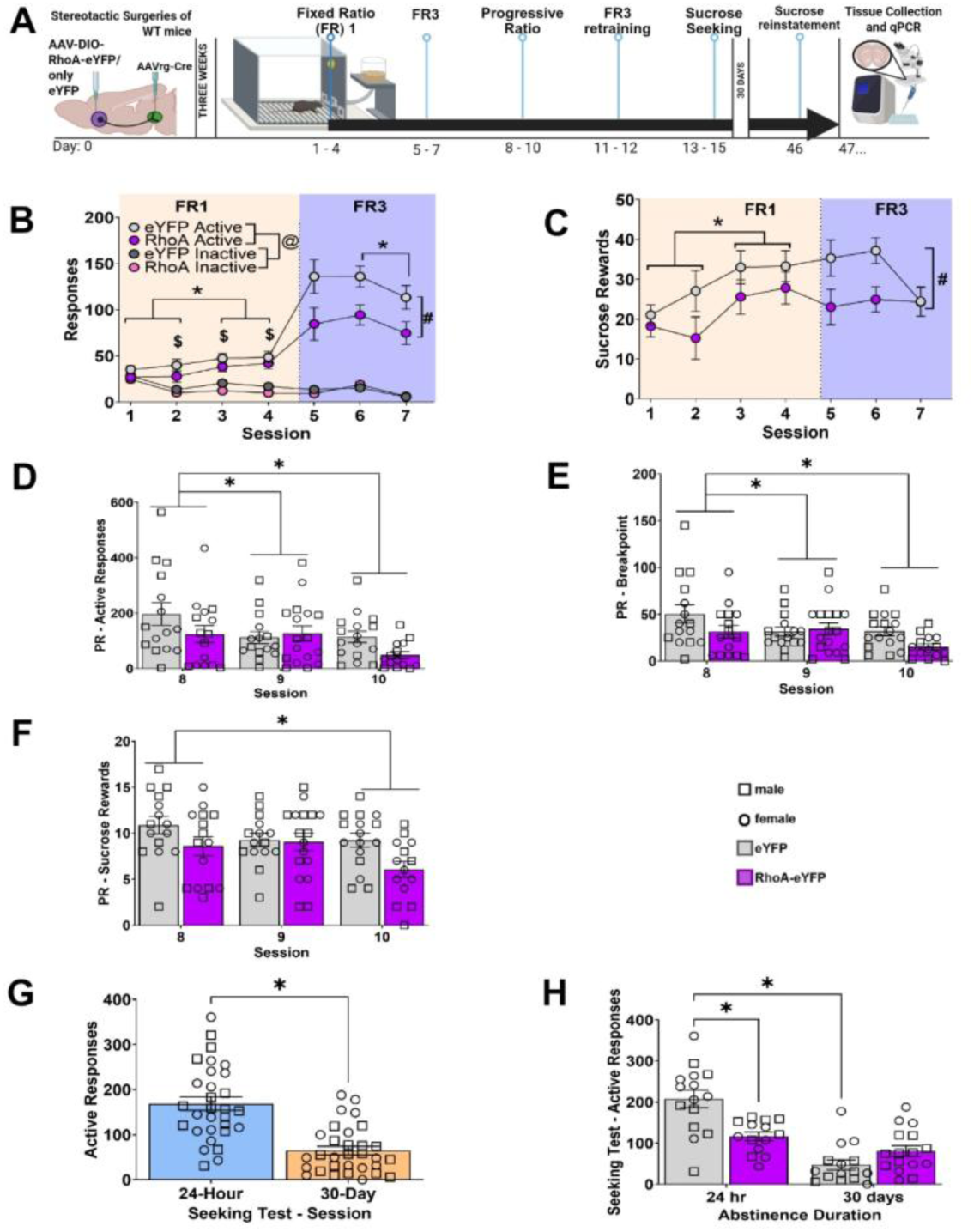
RhoA-OE in NAc-VTA D1-SPNs blunted sucrose operant responding. (A) Experimental timeline. AAV-Rg-Cre was injected into the VTA, and AAV-DIO-RhoA-eYFP or AAV-DIO-eYFP was injected into the NAc of wild-type mice. Following a 3-week viral-expression period, mice underwent fixed ratio 1 (FR1; days 1–4), FR3 (days 5–7), progressive ratio (PR; days 8–10), and FR3 retraining (days 11–12) for sucrose, followed by sucrose seeking (days 13–15) and, after 30 days of abstinence, sucrose reinstatement (day 46). Tissue was collected for RT-qPCR the next day (day 47). (B) Active and inactive nose-poke responses during FR1 (sessions 1–4, orange shading) and FR3 (sessions 5–7, blue shading) training. Mice responded more at the active than the inactive port (@), with greater active than inactive responding on sessions 2, 3, and 4 ($). Active responses were higher on sessions 3 and 4 than on sessions 1 and 2 (*) regardless of AAV treatment. During FR3, RhoA-OE mice made fewer active responses than eYFP controls (#), and responding decreased from session 6 to session 7 (*). (C) Sucrose rewards earned during FR1 and FR3 training. Rewards earned did not differ between RhoA-OE and eYFP mice during FR1, and all mice earned more rewards on sessions 3 and 4 than on sessions 1 and 2 (*). During FR3, RhoA-OE mice earned fewer rewards than eYFP controls (#). (D) Active responses decreased across PR sessions in both groups (*, session 8 vs. sessions 9 and 10). (E) Breakpoint was lower on sessions 9 and 10 than on session 8 in both groups (*). (F) The number of sucrose rewards earned was lower on session 10 than on session 8 in both groups (*). (G) Across all mice, active responding in the cue-induced sucrose seeking test was lower after 30 days of abstinence than after 24 hr of abstinence (*; blue bar: 24-hour, orange bar: 30-day). (H) RhoA-OE mice made fewer active responses than eYFP controls after 24 hr of abstinence, but not after 30 days (*). eYFP controls made fewer active responses after 30 days than after 24 hr of abstinence (*). (Squares: males, circles: females; gray bars: eYFP, purple bars: RhoA-eYFP; all figures represent mean ± SEM, statistical significance was assessed using p = 0.05, *p < 0.05, **p < 0.01, ***p < 0.001, and ****p < 0.0001; ns, not significant.)

### Sucrose self-administration under a fixed ratio schedule of reinforcement

To first test whether RhoA-OE in NAc-VTA projection neurons impacts sucrose operant behavior. Mice were tested for 4 sessions on an FR1 schedule of reinforcement. Analysis of active port responding (**Fig. 3B**) or sucrose reinforcers (**Fig. 3C**) showed no significant effect of AAV treatment (F(1,25) = 1.88, p = 0.183; F(1,24) = 1.72, p = 0.202, respectively), sex (F(1,21) = 2.63, p = 0.120; F(1,21) = 1.80, p = 0.200) or interaction between the two (F(2.44,61) = 0.23, p = 0.840; F(2.32,55.73) = 1.70, p = 0.200, respectively). However, a significant effect of port revealed that both eYFP and RhoA-OE groups increased responding and biasing for the active sucrose-paired port over the inactive port (**Fig. 3B**; F(1,69) = 25.13, p < 0.001). Furthermore, a significant port-by-day interaction (F(3,69) = 18.05, p < 0.001) and post-hoc comparisons showed greater responding at the active than inactive port by day 2 (Bonferroni, p ≤ 0.001). Thus, these findings suggest that eYFP and RhoA-OE mice showed comparable acquisition of the operant task, with both groups developing a preference for the sucrose-paired port over the inactive port by session two on an FR1 schedule.

Mice were subsequently tested under an FR3 schedule to determine whether RhoA-OE in NAc-VTA SPNs affected responding under a higher response requirement. Here, a significant effect of AAV treatment showed that RhoA-OE attenuated active responding across the three sessions (**Fig. 3B**; F(1,25) = 5.49, p = 0.027) but trended towards a decrease in the number of sucrose reinforcers earned (**Fig. 3C**; F(1,25) = 4.10, p = 0.055) relative to eYFP controls. Sex did not have an effect on active responding (F(1,25) = 0.043, p = 0.840) or sucrose (F(1,25) = 0.24, p = 0.630) reinforcement, and there was no sex by AAV treatment interaction for either measure (F(1,25) = 0.030, p = 0.870; F(1,25) = 0.001, p = 0.974, respectively). These findings indicate that RhoA-OE in NAc-VTA D1-SPNs was associated with a reduction in sucrose-directed responding under the FR3 schedule.

### Sucrose self-administration under a progressive ratio schedule of reinforcement

To assess whether RhoA-OE in NAc-VTA SPNs alters incentive motivation for sucrose, mice were transitioned to a PR schedule following FR3 training (**Fig. 3D-F**). Analysis of PR performance revealed a significant effect of AAV treatment on the number of sucrose reinforcers earned (F(1,23) = 4.70, p = 0.041), with RhoA-OE mice earning fewer sucrose reinforcers than eYFP controls (**Fig. 3F**). Although AAV treatment did not significantly affect active-port responding or breakpoint (F(1,23) = 2.60, p = 0.123; F(1,23) = 3.65, p = 0.07), RhoA-OE mice exhibited lower active-port responding than eYFP mice, corresponding to an approximately 40% reduction (**Fig. 3D**). There was also a significant effect of PR training day on active-port responding (F(1.60,36.70) = 11.02, p < 0.001), breakpoint (F(1.60,36.86) = 9.24, p = 0.001), and sucrose reinforcement (F(2.0,44.13) = 8.40, p < 0.001). Bonferroni-corrected pairwise comparisons indicated that active-port responding (p < 0.001), breakpoint (p = 0.030), and sucrose reinforcers earned (p < 0.001) decreased by third day of PR training (**Fig. 3D-F**). Together, these findings indicate that RhoA-OE in NAc-VTA SPNs reduces sucrose reinforcement under a PR schedule, consistent with reduced incentive motivation for sucrose.

### RhoA overexpression attenuates cue-induced sucrose seeking after acute abstinence from self-administration

Lastly, to determine whether RhoA-OE in NAc-VTA SPNs impacts relapse-like vulnerability following acute and prolonged abstinence, mice were tested for cue-induced sucrose seeking after 24 h and 30 days of presentation of the sucrose-associated cue but did not deliver sucrose pellets. Analysis of active-port responding revealed a significant interaction between abstinence duration and AAV treatment (F(1,27) = 18.28, p < 0.001). Post-hoc comparisons showed that RhoA-OE mice exhibited fewer active-port responses than eYFP mice following 24 h of abstinence (Bonferroni, p = 0.001), whereas this difference was not significant following 30 days of abstinence (p = 0.062; **Fig. 3H**). Additionally, eYFP mice exhibited a 78% reduction in active-port responding from 24 h to 30 days of abstinence (Bonferroni, p < 0.001), whereas the 31% reduction observed in RhoA-OE mice was not significant across abstinence duration (Bonferroni, p = 0.086; **Fig. 3H**). There was also a significant effect of abstinence duration (F(1,27) = 45.73, p < 0.001), with fewer active-port responses after 30 days compared with 24 h of abstinence (**Fig. 3G**). There was no effect AAV treatment (F(1,27) = 1.16, p = 0.08) or sex (F(1,27) = 3.32, p = 0.29) on cue-induced sucrose seeking. Together, these findings indicate that the effect of NAc RhoA-OE on cue-induced sucrose seeking differed between acute and prolonged abstinence, with RhoA-OE reducing cue-induced responding after 24 h but not 30 days of abstinence.

### CSDS and CWDS induced RhoA mRNA expression in NAc to VTA D1-SPNs in NAc

We first examined RhoA mRNA expression in NAc-VTA D1-SPNs using the experimental approach shown in **Fig. 4A**. To selectively label NAc-VTA SPNs, a retrograde AAV-Rg-Cre vector was injected into the VTA of wild-type mice. Following three weeks to allow for optimal viral expression, male mice underwent chronic social defeat stress and female mice underwent chronic witness defeat stress. Behavioral and molecular data were subsequently analyzed using one-way ANOVAs, with stress group social interactor (CTRL vs. LOW vs. HIGH) as a between-subject factor.

**Figure 4.**
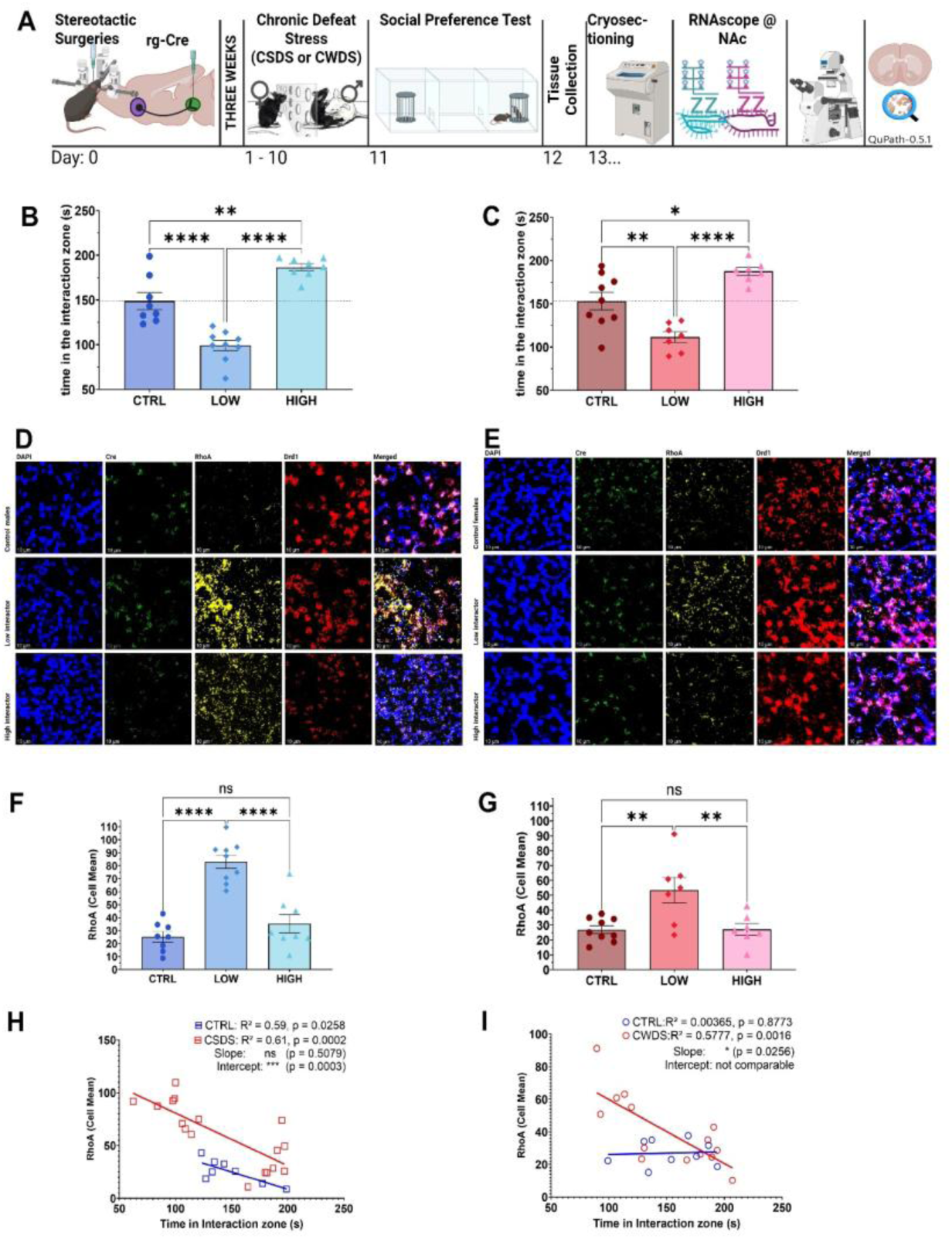
RhoA mRNA expression in NAc-VTA D1-SPNs was elevated in Low Social Interactors after CSDS and CWDS and was negatively correlated with social interaction. (A) Experimental timeline. AAV-Rg-Cre was injected into the VTA of wild-type mice. Following a 3-week viral-expression period, mice underwent chronic social defeat stress (CSDS; males) or chronic witness defeat stress (CWDS; females) on days 1–10, followed by the three-chamber social interaction test (3CSI) on day 11. Brains were collected on day 12 and cryosectioned (day 13) for multiplex RNAscope of the NAc, and images were analyzed using QuPath (v0.5.1). (B, C) In the 3CSI, stress-exposed mice were classified as Low Social Interactors (LOW) or High Social Interactors (HIGH) according to whether their interaction-zone time was below or above the mean of sex-matched control (CTRL) mice (dotted line). LOW males (B) and LOW females (C) spent less time in the interaction zone, whereas HIGH males and HIGH females spent more time in the interaction zone than CTRL and LOW mice. (D, E) Representative RNAscope images of the NAc from CTRL, LOW, and HIGH males following CSDS (D) and females following CWDS (E), showing DAPI (blue), Cre (green), RhoA (yellow), Drd1 (red), and the merged image. Scale bars, 10 µm. (F, G) Quantification of RhoA mRNA expression (cell mean) in the NAc of males (F) and females (G). RhoA mRNA expression was greater in LOW mice than in CTRL and HIGH mice, whereas it did not differ between CTRL and HIGH mice. (H, I) Simple linear regression of RhoA mRNA expression against interaction-zone time. RhoA mRNA expression was negatively correlated with interaction-zone time in CSDS-exposed males (H; red) and CWDS-exposed females (I; red). In CTRL mice (blue), RhoA mRNA expression was negatively correlated with interaction-zone time in males (H) but not in females (I). (All figures represent mean ± SEM, statistical significance was assessed using p = 0.05, *p < 0.05, **p < 0.01, ***p < 0.001, and ****p < 0.0001; ns, not significant.)

Using three-chamber social interaction test, stress-exposed mice were classified as Low Social Interactors (LOW) or High Social Interactors (HIGH), according to whether their interaction-zone time was below or above the mean value of sex-matched control mice. In males exposed to CSDS, a significant main effect of social interaction indicated that interaction zone time differed significantly among control (CTRL), LOW, and HIGH groups (**Fig. 4B**; F(2,22) = 43.41, p < 0.0001). Post-hoc comparisons revealed that LOW males spent significantly less time in the interaction zone (Tukey, CTRL vs. LOW: p < 0.0001), whereas HIGH males spent significantly more time in the interaction zone than CTRL and LOW males (Tukey, CTRL vs. effect of social interactor indicated that interaction zone time differed significantly among the three groups (**Fig. 4C**; one-way ANOVA, F(2,20) = 20.22, p < 0.0001). Post-hoc comparisons showed that LOW females spent significantly less time in the interaction zone than controls (Tukey, CTRL vs. LOW: p = 0.0039). However, HIGH females spent significantly more time in the interaction zone than control and LOW females (Tukey, CTRL vs. HIGH: p = 0.0172; LOW vs. HIGH: p < 0.0001). These findings confirm distinct social-interaction phenotypes following social defeat stress in males and females, with LOW animals exhibiting reduced social interaction and HIGH animals exhibiting enhanced social interaction relative to controls.

RNAscope images from male and female mice are shown in **Fig. 4D** and **4E**, respectively, and were quantified using QuPath (v0.5.1; Bankhead et al. 2017). RhoA mRNA expression (average number of puncta per cell) in VTA-projecting, Cre-positive, Drd1-positive NAc neurons differed significantly among control, LOW, and HIGH groups following CSDS in males (**Fig. 4F**; F (2, 22) = 31.86, p<0.0001). Post-hoc comparisons revealed significantly greater RhoA mRNA expression in LOW males compared to CTRL and HIGH males (Tukey, CTRL vs. LOW: p < 0.0001; LOW vs. HIGH: p < 0.0001), whereas RhoA expression did not differ between CTRL and HIGH males (Tukey, CTRL vs. HIGH: p = 0.4400). In females following CWDS, RhoA mRNA expression also differed significantly among CTRL, LOW, and HIGH groups (**Fig. 4G**; F (2, 20) = 8.184, p=0.0025). Post-hoc comparisons demonstrated significantly greater RhoA expression in LOW females compared to CTRL and HIGH females (Tukey, CTRL vs. LOW: p = 0.0043; LOW vs. HIGH: p = 0.0074), whereas RhoA expression did not differ between CTRL and HIGH females (Tukey, CTRL vs. HIGH: p = 0.9994).

Finally, we examined the relationship between RhoA mRNA expression and interaction-zone time. In males, RhoA expression was negatively correlated with interaction-zone time in both CTRL mice (**Fig. 4H**; R² = 0.59, p = 0.0258) and CSDS-exposed mice (R² = 0.61, p = 0.0002). In females, RhoA expression was negatively correlated with interaction-zone time in CWDS-exposed mice (**Fig. 4I**; R² = 0.5777, p = 0.0016), but not in control females (R² = 0.00365, p = 0.8773). Together, these findings indicate that increased RhoA mRNA expression in VTA-projecting D1-SPNs from NAc is associated with reduced social interaction following social defeat stress.

## Discussion

Stress research field has identified the projection-defined subgroups of NAc-D1-SPNs projecting to distinct downstream targets (Kupchik et al. 2015; Baimel et al. 2019; Pardo-Garcia et al. 2019; Enriquez-Traba et al. 2025) and is gradually identifying their molecular identity and functional roles. Whether RhoA signaling operates uniformly across these subgroups remains unexplored. Our findings identify RhoA signaling within NAc-VTA D1-SPNs as a critical molecular mechanism linking chronic stress to maladaptive behavioral outcomes. First, projection-specific RhoA-OE in NAC-VTA SPNs was sufficient to induce a negative affective phenotype, characterized by increased anxiety-like behavior, social avoidance, reduced grooming, and reduced food approach, without significantly affecting locomotor activity. Second, RhoA-OE significantly reduced reward-directed behavior. Third, chronic stress increased RhoA mRNA expression specifically in NAc-VTA. Finally, stress susceptibility was associated with elevated RhoA expression within the NAc: higher levels of RhoA expression were correlated with greater social avoidance.

Reduced social preference and reduced grooming reproduce what is observed when RhoA is overexpressed across all D1-SPNs (Fox and Lobo 2019; Fox et al. 2020a), and align with the studies showing that inhibiting RhoA with Rhosin into NAc during defeat stress promotes resilience (Francis et al. 2019). Although, D1-SPN RhoA-OE left elevated plus maze open-arm time and open field center time unchanged in both sexes (Fox et al. 2020a), whereas restricting overexpression to VTA-projecting neurons produced avoidance of both, together with delayed feeding in a novel arena. Therefore, negative affective-like avoidance may be carried disproportionately by the NAc-VTA pathway and obscured when all D1-SPNs have RhoA-OE. Total distance travelled in both control (eYFP) and RhoA-OE (RhoA-eYFP) groups does not differ in case of habituation phase of three-chamber social interaction test and for open-field test shows RhoA-OE does not impair the locomotion of the mice. However, the distance traveled in the elevated zero maze for RhoA-OE groups was significantly less compared to the control; this could possibly stem from more freezing episodes at the junction of the closed and open arms for the RhoA-OE group compared to the control group, suggesting an anxiety-like phenotype.

Additionally, the present findings show that RhoA signaling in NAc-VTA SPNs selectively regulates motivation for sucrose reward. RhoA-OE did not impair acquisition of the sucrose operant task, but it reduced responding when greater effort was required and attenuated cue-induced sucrose seeking following acute abstinence. This suggests a decreased willingness to work for and seek reward. Previous work has shown that RhoA-OE in NAc D1-SPNs decreases sucrose preference and promotes a stress-susceptible phenotype characterized by social avoidance (Fox et al. 2020a). Thus, enhanced RhoA signaling may contribute to reduced reward-directed behavior by altering affective state. Consistent with this interpretation, the present findings showed RhoA-OE mice exhibited reduced grooming in the sucrose splash test, a behavioral measure commonly interpreted as reduced self-care and depressive-like behavior (Yalcin et al. 2008). Manipulations of the RhoA/ROCK pathway have been shown to influence response to both natural and drug rewards (DePoy et al. 2013; Swanson et al. 2017). Together, these observations suggest that increased RhoA signaling within NAc-VTA SPNs reduces motivation for sucrose and promote negative affective-like states that compete with or suppress appetitive behavior (DePoy et al. 2013; Cahill et al. 2018; Francis et al. 2019; Fox and Lobo 2019; Fox et al. 2020a).

The RNAscope experiment shows that chronic social stress engages RhoA in NAc-VTA D1-SPNs. Retrograde labeling confirmed that approximately 90% of VTA-projecting NAc neurons express Drd1, so RhoA induction measured in this population can be attributed primarily to D1-SPNs, consistent with our previous studies in male CSDS mice. Both CSDS in males and CWDS in females increased RhoA mRNA in these NAc-VTA D1-SPNs, extending stress-induced RhoA D1-SPN induction to a defined output pathway (Francis et al. 2019; Fox et al. 2020a). The negative correlation between RhoA expression and interaction-zone time confirms the role of RhoA in chronic stress-mediated negatively affective behavior. This finding is consistent with the previously described role of RhoA in the whole NAc (Fox and Lobo 2019; Fox et al. 2020a). The same induction in both sexes, together with comparable behavioral effects of RhoA-OE, suggests a shared instead of sex-specific contributor to stress susceptibility. Accordingly, vicarious defeat acts through psychological stimuli rather than a physical stressor (Warren et al. 2013; Sial et al. 2016; Iñiguez et al. 2018).

Three mechanisms could link elevated RhoA to negative affective-like behaviors.

*The first is structural*. RhoA acts through ROCK on the actin cytoskeleton to regulate spine morphology and dendritic remodeling (Penzes and Jones 2008; Penzes and Rafalovich 2012; Martino et al. 2013; Newell-Litwa et al. 2015; Konietzny et al. 2017; Mulherkar et al. 2017; Mulherkar and Tolias 2020). Constitutively active RhoA reduces dendritic complexity in D1-SPNs, and ROCK inhibition after CSDS restores it (Francis et al. 2019; Fox et al. 2020a). A smaller arbor would reduce the surface available for cortical and limbic glutamatergic input. Thus, a smaller arbor would lower recruitment of these neurons during behavior. The same actin-dependent plasticity underlies structural adaptation to drugs of abuse (Mulholland and Chandler 2007; Dietz et al. 2009; Russo et al. 2010; Eisinger et al. 2020).

*The second is physiological*. Rhosin blocks CSDS-induced D1-SPN hyperexcitability and the accompanying loss of excitatory drive (Francis et al. 2015, 2019; Francis and Lobo 2017). The reduced excitatory transmission onto D1-SPNs accompanies anhedonic phenotypes (Lim et al. 2012; Lucantonio et al. 2025). These D1-SPNs are GABAergic and inhibit the inhibitory VTA neurons that regulate dopamine neurons. As a result, VTA dopamine neurons face more inhibition and secrete less dopamine (Lim et al. 2012; Soares-Cunha et al. 2020; Zhou et al. 2022). Therefore, negative affective-like behavior manifests.

*The third is transcriptional*. Egr3 binds the RhoA promoter and is required for stress-induced dendritic atrophy and social avoidance in D1-SPNs (Francis et al. 2017, 2019). BDNF-TrkB signaling in D1-SPNs regulates their plasticity and reward-related output (Stern 2010; Smith et al. 2013; Mason et al. 2013; Francis and Lobo 2017; Pagliusi et al. 2022). Therefore, an Egr3-RhoA axis modulated by VTA-derived BDNF can be a mechanism (Berton et al. 2006; Krishnan et al. 2007).

Clinical interest in RhoA-ROCK signaling rests on the availability of inhibitors that reverse stress-induced behavioral and structural deficits in rodents (Francis et al. 2019; Fox et al. 2020a). Our results indicate the behavioral consequences of manipulating RhoA depend on the circuit engaged, so the net effect of a systemically delivered ROCK inhibitor could impact multiple pathways within the NAc and across the brain, thus deeper investigation into RhoA mechanisms across reward and motivation brain circuitry is warranted. Effort-based motivational deficits of this kind map onto anhedonia and amotivation, symptoms poorly served by existing antidepressants (Al-harbi 2012; Dodd et al. 2021; Touloumis 2021; Havlik et al. 2024; Siyoum et al. 2024; Fiorillo et al. 2025), and their appearance in both sexes strengthens the case for RhoA mechanism within this NAc-VTA pathway, as well as other brain circuits..

This study has limitations that define the scope of its conclusions. CSDS was performed in males and CWDS in females, so sex-based comparison of stress-induced RhoA expression (Fig. 4) is not possible; the effects of RhoA-OE, in contrast, were assessed in both sexes in stress naïve conditions.

Collectively our studies uncover the behavioral consequences of RhoA-OE in NAc-VTA SPNs that are primarily D1-SPNs. By localizing RhoA-dependent behavioral effects to a defined output pathway, and showing that effort-based measures of motivation as well as affective and exploratory behaviors are sensitive to this molecular alterations, this work provides a framework for evaluating whether circuit-informed modulation of RhoA-ROCK signaling can address anhedonia, amotivation, and anxiety-like outcomes. Providing an early step to potential translational utility of RhoA mechanisms in the future.

## Funding

This work was supported by R01MH106500 and R01DA038613 to MKL and F32DA066191 to SNS. All viral vectors were generated within the Viral Vector Core at the University of Maryland School of Medicine.

## Competing interests

The authors declare that they have no competing interests.

## Data availability

All data supporting the conclusions of this article are presented within the article. Raw data, viral construct maps, and analysis scripts are available from the corresponding author upon reasonable request.

## Ethics approval

All animal procedures were approved by the Institutional Animal Care and Use Committee (IACUC) at the University of Maryland School of Medicine and conformed to the National Institutes of Health Guide for the Care and Use of Laboratory Animals.

## Notes

### Competing Interest Statement

The authors have declared no competing interest.

## Reference

Al-harbi (2012) Treatment-resistant depression: therapeutic trends, challenges, and future directions. PPA 369. 10.2147/PPA.S29716

Baimel C, McGarry LM, Carter AG (2019) The projection targets of medium spiny neurons govern cocaine-evoked synaptic plasticity in the nucleus accumbens. Cell Rep 28:2256–2263.e3. 10.1016/j.celrep.2019.07.074

Bankhead P, Loughrey MB, Fernández JA, et al (2017) QuPath: open source software for digital pathology image analysis. Sci Rep 7:16878. 10.1038/s41598-017-17204-5

Berridge KC, Kringelbach ML (2015) Pleasure Systems in the Brain. Neuron 86:646–664. 10.1016/j.neuron.2015.02.018

Berton O, McClung CA, DiLeone RJ, et al (2006) Essential Role of BDNF in the Mesolimbic Dopamine Pathway in Social Defeat Stress. Science 311:864–868. 10.1126/science.1120972

Bocklisch C, Pascoli V, Wong JCY, et al (2013) Cocaine disinhibits dopamine neurons by potentiation of GABA transmission in the ventral tegmental area. Science 341:1521–1525. 10.1126/science.1237059

Bodas D, Demirci Ş, Balcou M, et al (2026) A state-dependent neural circuit resolves approach– avoidance conflicts

Brown JS (1948) Gradients of approach and avoidance responses and their relation to level of motivation. J Comp Physiol Psychol 41:450–465. 10.1037/h0055463

Burek DJ, Carlezon WA (2026) Stress-induced adaptations in nucleus accumbens dopamine D1 receptor-expressing cells correspond to social avoidance behavior in male mice. IBRO Neurosci Rep 21:335–341. 10.1016/j.ibneur.2026.07.009

Cahill ME, Browne CJ, Wang J, et al (2018) Withdrawal from repeated morphine administration augments expression of the RhoA network in the nucleus accumbens to control synaptic structure. J Neurochem 147:84–98. 10.1111/jnc.14563

Calarco CA, Fox ME, Van Terheyden S, et al (2021) Mitochondria-Related Nuclear Gene Expression in the Nucleus Accumbens and Blood Mitochondrial Copy Number After Developmental Fentanyl Exposure in Adolescent Male and Female C57BL/6 Mice. Front Psychiatry 12:737389. 10.3389/fpsyt.2021.737389

Carlezon WA, Thomas MJ (2009) Biological substrates of reward and aversion: A nucleus accumbens activity hypothesis. Neuropharmacology 56:122–132. 10.1016/j.neuropharm.2008.06.075

Carter CA, Pudipeddi SS, Llorach P, et al (2025) Divergent Roles of Nucleus Accumbens D1-and D2-MSNs in Regulating Hedonic Feeding

Chandra R, Francis TC, Nam H, et al (2017) Reduced Slc6a15 in Nucleus Accumbens D2-Neurons Underlies Stress Susceptibility. J Neurosci 37:6527–6538. 10.1523/JNEUROSCI.3250-16.2017

Cléry-Melin M-L, Jollant F, Gorwood P (2019) Reward systems and cognitions in Major Depressive Disorder. CNS Spectr 24:64–77. 10.1017/S1092852918001335

Correia R, Coimbra B, Domingues AV, et al (2023) Involvement of nucleus accumbens D2– medium spiny neurons projecting to the ventral pallidum in anxiety-like behaviour. J Psychiatry Neurosci 48:E267–E284. 10.1503/jpn.220111

Daniels A, Wellan SA, Beck A, et al (2025) Anhedonia relates to reduced striatal reward anticipation in depression but not in schizophrenia or bipolar disorder: A transdiagnostic study. Cogn Affect Behav Neurosci 25:501–514. 10.3758/s13415-024-01261-1

DePoy LM, Noble B, Allen AG, Gourley SL (2013) Developmentally divergent effects of Rho-kinase inhibition on cocaine-and BDNF-induced behavioral plasticity. Behav Brain Res 243:171–175. 10.1016/j.bbr.2013.01.004

Deussing JM (2006) Animal models of depression. Drug Discov Today Dis Models 3:375–383. 10.1016/j.ddmod.2006.11.003

Dietz DM, Dietz KC, Nestler EJ, Russo SJ (2009) Molecular Mechanisms of Psychostimulant-induced Structural Plasticity. Pharmacopsychiatry 42:S69–S78. 10.1055/s-0029-1202847

Dodd S, Bauer M, Carvalho AF, et al (2021) A clinical approach to treatment resistance in depressed patients: What to do when the usual treatments don’t work well enough? World J Biol Psychiatry 22:483–494. 10.1080/15622975.2020.1851052

Du W, Jia Y, Hu C (2026) Global, Regional, and National Burden of Major Depressive Disorder From 1990 to 2021: Analysis of Data From GBD 2021. Ann Clin Psychiatry 10401237261473586. 10.1177/10401237261473586

Edwards NJ, Tejeda HA, Pignatelli M, et al (2017) Circuit specificity in the inhibitory architecture of the VTA regulates cocaine-induced behavior. Nat Neurosci 20:438–448. 10.1038/nn.4482

Eisinger KRT, Chapp AD, Swanson SP, et al (2020) Caveolin-1 regulates medium spiny neuron structural and functional plasticity. Psychopharmacology (Berl) 237:2673–2684. 10.1007/s00213-020-05564-2

Enriquez-Traba J, Arenivar M, Yarur-Castillo HE, et al (2025) Dissociable control of motivation and reinforcement by distinct ventral striatal dopamine receptors. Nat Neurosci 28:105–121. 10.1038/s41593-024-01819-9

Faust TW, Mohebi A, Berke JD (2025) Reward expectation and receipt differentially modulate the spiking of accumbens D1+ and D2+ neurons. Curr Biol 35:1285–1297.e3. 10.1016/j.cub.2025.02.007

Fiorillo A, Demyttenaere K, Martiadis V, Martinotti G (2025) Editorial: Treatment resistant depression (TRD): epidemiology, clinic, burden and treatment. Front Psychiatry 16:1588902. 10.3389/fpsyt.2025.1588902

Flores RJ, Jaime-Lara RB, Arenivar MA, et al (2026) High-fat diet exposure, internal state, and reward value influence decision making during motivational conflict. Neurobiol Stress 44:100844. 10.1016/j.ynstr.2026.100844

Fox ME, Chandra R, Menken MS, et al (2020a) Dendritic remodeling of D1 neurons by RhoA/Rho-kinase mediates depression-like behavior. Mol Psychiatry 25:1022–1034. 10.1038/s41380-018-0211-5

Fox ME, Figueiredo A, Menken MS, Lobo MK (2020b) Dendritic spine density is increased on nucleus accumbens D2 neurons after chronic social defeat. Sci Rep 10:12393. 10.1038/s41598-020-69339-7

Fox ME, Lobo MK (2019) The molecular and cellular mechanisms of depression: a focus on reward circuitry. Mol Psychiatry 24:1798–1815. 10.1038/s41380-019-0415-3

Francis TC, Chandra R, Friend DM, et al (2015) Nucleus Accumbens Medium Spiny Neuron Subtypes Mediate Depression-Related Outcomes to Social Defeat Stress. Biol Psychiatry 77:212–222. 10.1016/j.biopsych.2014.07.021

Francis TC, Chandra R, Gaynor A, et al (2017) Molecular basis of dendritic atrophy and activity in stress susceptibility. Mol Psychiatry 22:1512–1519. 10.1038/mp.2017.178

Francis TC, Gaynor A, Chandra R, et al (2019) The Selective RhoA Inhibitor Rhosin Promotes Stress Resiliency Through Enhancing D1-Medium Spiny Neuron Plasticity and Reducing Hyperexcitability. Biol Psychiatry 85:1001–1010. 10.1016/j.biopsych.2019.02.007

Francis TC, Lobo MK (2017) Emerging Role for Nucleus Accumbens Medium Spiny Neuron Subtypes in Depression. Biol Psychiatry 81:645–653. 10.1016/j.biopsych.2016.09.007

Franco D, Wulff AB, Lobo MK, Fox ME (2022) Chronic Physical and Vicarious Psychosocial Stress Alter Fentanyl Consumption and Nucleus Accumbens Rho GTPases in Male and Female C57BL/6 Mice. Front Behav Neurosci 16:821080. 10.3389/fnbeh.2022.821080

Golden SA, Christoffel DJ, Heshmati M, et al (2013) Epigenetic regulation of RAC1 induces synaptic remodeling in stress disorders and depression. Nat Med 19:337–344. 10.1038/nm.3090

Golden SA, Covington HE, Berton O, Russo SJ (2011) A standardized protocol for repeated social defeat stress in mice. Nat Protoc 6:1183–1191. 10.1038/nprot.2011.361

Grubbs FE (1969) Procedures for detecting outlying observations in samples. Technometrics 11:1–21. 10.1080/00401706.1969.10490657

Grubbs FE (1950) Sample Criteria for Testing Outlying Observations. Ann Math Stat 21:27–58. 10.1214/aoms/1177729885

Gutierrez R, Lobo MK, Zhang F, de Lecea L (2011) Neural integration of reward, arousal, and feeding: Recruitment of VTA, lateral hypothalamus, and ventral striatal neurons. IUBMB Life 63:824–830. 10.1002/iub.539

Halahakoon DC, Kieslich K, O’Driscoll C, et al (2020) Reward-Processing Behavior in Depressed Participants Relative to Healthy Volunteers: A Systematic Review and Meta-analysis. JAMA Psychiatry 77:1286. 10.1001/jamapsychiatry.2020.2139

Hamel L, Thangarasa T, Samadi O, Ito R (2017) Caudal Nucleus Accumbens Core Is Critical in the Regulation of Cue-Elicited Approach-Avoidance Decisions. eneuro 4:ENEURO.0330-16.2017. 10.1523/ENEURO.0330-16.2017

Hammen C, Kim EY, Eberhart NK, Brennan PA (2009) Chronic and acute stress and the prediction of major depression in women. Depress Anxiety 26:718–723. 10.1002/da.20571

Havlik JL, Wahid S, Teopiz KM, et al (2024) Recent Advances in the Treatment of Treatment-Resistant Depression: A Narrative Review of Literature Published from 2018 to 2023. Curr Psychiatry Rep 26:176–213. 10.1007/s11920-024-01494-4

Ilango A, Lobo MK (2015) Shining light on motivation, emotion, and memory processes. Front Behav Neurosci 9:. 10.3389/fnbeh.2015.00001

Illescas-Huerta E, Hernández-Ortiz E, Sotres-Bayon F (2026) Top-down corticostriatal control of adaptive restraint during motivational conflict. Sci Adv 12:eaeg2327. 10.1126/sciadv.aeg2327

Iñiguez SD, Flores-Ramirez FJ, Riggs LM, et al (2018) Vicarious Social Defeat Stress Induces Depression-Related Outcomes in Female Mice. Biol Psychiatry 83:9–17. 10.1016/j.biopsych.2017.07.014

Kai N, Nishizawa K, Tsutsui Y, et al (2015) Differential roles of dopamine D1 and D2 receptor-containing neurons of the nucleus accumbens shell in behavioral sensitization. J Neurochem 135:1232–1241. 10.1111/jnc.13380

Khayat A, Yaka R (2024) Activation of nucleus accumbens projections to the ventral tegmental area alters molecular signaling and neurotransmission in the reward system. Front Mol Neurosci 17:1271654. 10.3389/fnmol.2024.1271654

Kokras N, Baltas D, Theocharis F, Dalla C (2017) Kinoscope: An Open-Source Computer Program for Behavioral Pharmacologists. Front Behav Neurosci 11:88. 10.3389/fnbeh.2017.00088

Konietzny A, Bär J, Mikhaylova M (2017) Dendritic Actin Cytoskeleton: Structure, Functions, and Regulations. Front Cell Neurosci 11:147. 10.3389/fncel.2017.00147

Kraeuter A-K, Guest PC, Sarnyai Z (2019) The Open Field Test for Measuring Locomotor Activity and Anxiety-Like Behavior. In: Guest PC (ed) Pre-Clinical Models. Springer New York, New York, NY, pp 99–103

Krishnan V, Han M-H, Graham DL, et al (2007) Molecular Adaptations Underlying Susceptibility and Resistance to Social Defeat in Brain Reward Regions. Cell 131:391–404. 10.1016/j.cell.2007.09.018

Krishnan V, Nestler EJ (2011) Animal Models of Depression: Molecular Perspectives. In: Hagan JJ (ed) Molecular and Functional Models in Neuropsychiatry. Springer Berlin Heidelberg, Berlin, Heidelberg, pp 121–147

Kumar G, Franco D, Basu M, et al (2026) Nucleus accumbens neuron subtype translatome signatures in socially stressed females. Neuropsychopharmacology 51:2026–2036. 10.1038/s41386-026-02414-1

Kupchik YM, Brown RM, Heinsbroek JA, et al (2015) Coding the direct/indirect pathways by D1 and D2 receptors is not valid for accumbens projections. Nat Neurosci 18:1230–1232. 10.1038/nn.4068

Lang PJ, Bradley MM (2010) Emotion and the motivational brain. Biol Psychol 84:437–450. 10.1016/j.biopsycho.2009.10.007

LeGates TA, Kvarta MD, Tooley JR, et al (2018) Reward behaviour is regulated by the strength of hippocampus–nucleus accumbens synapses. Nature 564:258–262. 10.1038/s41586-018-0740-8

Lim BK, Huang KW, Grueter BA, et al (2012) Anhedonia requires MC4R-mediated synaptic adaptations in nucleus accumbens. Nature 487:183–189. 10.1038/nature11160

Livak KJ, Schmittgen TD (2001) Analysis of relative gene expression data using real-time quantitative PCR and the 2−ΔΔCT method. Methods 25:402–408. 10.1006/meth.2001.1262

Lobo MK, Nestler EJ (2011) The Striatal Balancing Act in Drug Addiction: Distinct Roles of Direct and Indirect Pathway Medium Spiny Neurons. Front Neuroanat 5:. 10.3389/fnana.2011.00041

Lobo MK, Zaman S, Damez-Werno DM, et al (2013) FosB Induction in Striatal Medium Spiny Neuron Subtypes in Response to Chronic Pharmacological, Emotional, and Optogenetic Stimuli. J Neurosci 33:18381–18395. 10.1523/JNEUROSCI.1875-13.2013

Lucantonio F, Roeglin J, Li S, et al (2025) Ketamine rescues anhedonia by cell-type-and input-specific adaptations in the nucleus accumbens. Neuron 113:1398–1412.e4. 10.1016/j.neuron.2025.02.021

Malykhin N, Serrano J, Pietrasik W, Hegadoren K (2025) Effects of duration and intensity of psychological stressors on mental health outcomes. J Psychiatr Res 187:211–222. 10.1016/j.jpsychires.2025.05.012

Martino A, Ettorre M, Musilli M, et al (2013) Rho GTPase-dependent plasticity of dendritic spines in the adult brain. Front Cell Neurosci 7:. 10.3389/fncel.2013.00062

Mason BL, Lobo MK, Parada LF, Lutter M (2013) Trk B signaling in dopamine 1 receptor neurons regulates food intake and body weight: Trk B in D1R-Neurons Affects Body Weight. Obesity 21:2372–2376. 10.1002/oby.20382

McEwen BS (2008) Central effects of stress hormones in health and disease: Understanding the protective and damaging effects of stress and stress mediators. Eur J Pharmacol 583:174–185. 10.1016/j.ejphar.2007.11.071

McEwen BS (2012) Brain on stress: How the social environment gets under the skin. Proc Natl Acad Sci 109:17180–17185. 10.1073/pnas.1121254109

McEwen BS (2017) Neurobiological and Systemic Effects of Chronic Stress. Chronic Stress 1:2470547017692328. 10.1177/2470547017692328

McEwen BS (2007) Physiology and neurobiology of stress and adaptation: central role of the brain. Physiol Rev 87:873–904. 10.1152/physrev.00041.2006

McEwen BS, Gianaros PJ (2010) Central role of the brain in stress and adaptation: Links to socioeconomic status, health, and disease. Ann N Y Acad Sci 1186:190–222. 10.1111/j.1749-6632.2009.05331.x

McGonagle KA, Kessler RC (1990) Chronic stress, acute stress, and depressive symptoms. Am J Community Psychol 18:681–706. 10.1007/BF00931237

Mulherkar S, Firozi K, Huang W, et al (2017) RhoA-ROCK Inhibition Reverses Synaptic Remodeling and Motor and Cognitive Deficits Caused by Traumatic Brain Injury. Sci Rep 7:10689. 10.1038/s41598-017-11113-3

Mulherkar S, Tolias KF (2020) RhoA-ROCK Signaling as a Therapeutic Target in Traumatic Brain Injury. Cells 9:245. 10.3390/cells9010245

Mulholland PJ, Chandler LJ (2007) The Thorny Side of Addiction: Adaptive Plasticity and Dendritic Spines. Sci World J 7:9–21. 10.1100/tsw.2007.247

Murray CJ, Lopez AD (1997) Global mortality, disability, and the contribution of risk factors: Global Burden of Disease Study. The Lancet 349:1436–1442. 10.1016/S0140-6736(96)07495-8

Nestler EJ, Lüscher C (2019) The Molecular Basis of Drug Addiction: Linking Epigenetic to Synaptic and Circuit Mechanisms. Neuron 102:48–59. 10.1016/j.neuron.2019.01.016

Newell-Litwa KA, Badoual M, Asmussen H, et al (2015) ROCK1 and 2 differentially regulate actomyosin organization to drive cell and synaptic polarity. J Cell Biol 210:225–242. 10.1083/jcb.201504046

Pagliusi M, Franco D, Cole S, et al (2022) The BDNF-TrkB Pathway Acts Through Nucleus Accumbens D2 Expressing Neurons to Mediate Stress Susceptible Outcomes. Front Psychiatry 13:854494. 10.3389/fpsyt.2022.854494

Pardo-Garcia TR, Garcia-Keller C, Penaloza T, et al (2019) Ventral pallidum is the primary target for accumbens D1 projections driving cocaine seeking. J Neurosci 39:2041–2051. 10.1523/JNEUROSCI.2822-18.2018

Penzes P, Jones KA (2008) Dendritic spine dynamics – a key role for kalirin-7. Trends Neurosci 31:419–427. 10.1016/j.tins.2008.06.001

Penzes P, Rafalovich I (2012) Regulation of the Actin Cytoskeleton in Dendritic Spines. In: Kreutz MR, Sala C (eds) Synaptic Plasticity. Springer Vienna, Vienna, pp 81–95

Polter AM, Barcomb K, Tsuda AC, Kauer JA (2018) Synaptic function and plasticity in identified inhibitory inputs onto VTA dopamine neurons. Eur J Neurosci 47:1208–1218. 10.1111/ejn.13879

Qi G, Zhang P, Li T, et al (2022) NAc-VTA circuit underlies emotional stress-induced anxiety-like behavior in the three-chamber vicarious social defeat stress mouse model. Nat Commun 13:577. 10.1038/s41467-022-28190-2

Qin K, Wei Q (2026) Global and regional burden of major depressive disorder and associated risk factors: analysis from the Global Burden of disease study, 1990–2021. Eur Arch Psychiatry Clin Neurosci 276:2509–2520. 10.1007/s00406-026-02265-3

Richardson NR, Roberts DCS (1996) Progressive ratio schedules in drug self-administration studies in rats: a method to evaluate reinforcing efficacy. J Neurosci Methods 66:1–11. 10.1016/0165-0270(95)00153-0

Russo SJ, Dietz DM, Dumitriu D, et al (2010) The addicted synapse: mechanisms of synaptic and structural plasticity in nucleus accumbens. Trends Neurosci 33:267–276. 10.1016/j.tins.2010.02.002

Russo SJ, Nestler EJ (2013) The brain reward circuitry in mood disorders. Nat Rev Neurosci 14:609–625. 10.1038/nrn3381

Samuels BA, Hen R (2011) Novelty-Suppressed Feeding in the Mouse. In: Gould TD (ed) Mood and Anxiety Related Phenotypes in Mice. Humana Press, Totowa, NJ, pp 107–121

Santarelli L, Saxe M, Gross C, et al (2003) Requirement of hippocampal neurogenesis for the behavioral effects of antidepressants. Science 301:805–809. 10.1126/science.1083328

Seibenhener ML, Wooten MC (2015) Use of the Open Field Maze to Measure Locomotor and Anxiety-like Behavior in Mice. J Vis Exp 52434. 10.3791/52434

Shepherd JK, Grewal SS, Fletcher A, et al (1994) Behavioural and pharmacological characterisation of the elevated “zero-maze” as an animal model of anxiety. Psychopharmacology (Berl) 116:56–64. 10.1007/BF02244871

Sial OK, Warren BL, Alcantara LF, et al (2016) Vicarious social defeat stress: Bridging the gap between physical and emotional stress. J Neurosci Methods 258:94–103. 10.1016/j.jneumeth.2015.10.012

Siemonsmeier G, Iyer ES, Schuler H, et al (2026) Nucleus accumbens medial shell D1 and D2 medium spiny neurons signal the valence of conditioned stimuli in a sex-dependent manner. Neuropsychopharmacology. 10.1038/s41386-026-02460-9

Simmons DV, Petko AK, Paladini CA (2017) Differential expression of long-term potentiation among identified inhibitory inputs to dopamine neurons. J Neurophysiol 118:1998–2008. 10.1152/jn.00270.2017

Siyoum M, Kibrom E, Fanta T, et al (2024) Prevalence of treatment-resistant depression and associated factors among major depressive disorder follow-up patients at Saint Amanuel Mental Specialised Hospital in Ethiopia: a cross-sectional study. BMJ Open 14:e087006. 10.1136/bmjopen-2024-087006

Smith RJ, Lobo MK, Spencer S, Kalivas PW (2013) Cocaine-induced adaptations in D1 and D2 accumbens projection neurons (a dichotomy not necessarily synonymous with direct and indirect pathways). Curr Opin Neurobiol 23:546–552. 10.1016/j.conb.2013.01.026

Soares-Cunha C, De Vasconcelos NAP, Coimbra B, et al (2020) Nucleus accumbens medium spiny neurons subtypes signal both reward and aversion. Mol Psychiatry 25:3241–3255. 10.1038/s41380-019-0484-3

Stern PR (2010) BDNF, Dopamine, and Cocaine Reward. Sci Signal 3:. 10.1126/scisignal.3144ec326

Strekalova T, Svirin E, Gorlova A, et al (2023) Resilience and Vulnerability to Stress-Induced Anhedonia: Unveiling Brain Gene Expression and Mitochondrial Dynamics in a Mouse Chronic Stress Depression Model. Biomolecules 13:1782. 10.3390/biom13121782

Swanson AM, DePoy LM, Gourley SL (2017) Inhibiting Rho kinase promotes goal-directed decision making and blocks habitual responding for cocaine. Nat Commun 8:1861. 10.1038/s41467-017-01915-4

Touloumis C (2021) The burden and the challenge of treatment-resistant depression. Psychiatriki 32:11–14. 10.22365/jpsych.2021.046

Treadway MT, Bossaller NA, Shelton RC, Zald DH (2012) Effort-based decision-making in major depressive disorder: A translational model of motivational anhedonia. J Abnorm Psychol 121:553–558. 10.1037/a0028813

Treadway MT, Zald DH (2011) Reconsidering anhedonia in depression: lessons from translational neuroscience. Neurosci Biobehav Rev 35:537–555. 10.1016/j.neubiorev.2010.06.006

Turner RJ, Beiser M (1990) Major Depression and Depressive Symptomatology among the Physically Disabled: Assessing the Role of Chronic Stress. J Nerv Ment Dis 178:343–350. 10.1097/00005053-199006000-00001

Volman SF, Lammel S, Margolis EB, et al (2013) New Insights into the Specificity and Plasticity of Reward and Aversion Encoding in the Mesolimbic System. J Neurosci 33:17569–17576. 10.1523/JNEUROSCI.3250-13.2013

Von Mücke-Heim I-A, Urbina-Treviño L, Bordes J, et al (2023) Introducing a depression-like syndrome for translational neuropsychiatry: a plea for taxonomical validity and improved comparability between humans and mice. Mol Psychiatry 28:329–340. 10.1038/s41380-022-01762-w

Von Muecke-Heim I-A, Ries C, Urbina L, Deussing JM (2021) P2X7R antagonists in chronic stress-based depression models: a review. Eur Arch Psychiatry Clin Neurosci 271:1343– 1358. 10.1007/s00406-021-01306-3

Wang F, Flanagan J, Su N, et al (2012) RNAscope: a novel in situ RNA analysis platform for formalin-fixed, paraffin-embedded tissues. J Mol Diagn 14:22–29. 10.1016/j.jmoldx.2011.08.002

Wang F, Rong L, Luo L, et al (2024) Associations Between Psychological Stress and the Risk of First Onset of Major Depression Disorder: Results from a Longitudinal Study in 6,985 Chinese First-Year Students. Psychol Res Behav Manag Volume 17:3585–3593. 10.2147/PRBM.S482482

Warren BL, Vialou VF, Iñiguez SD, et al (2013) Neurobiological Sequelae of Witnessing Stressful Events in Adult Mice. Biol Psychiatry 73:7–14. 10.1016/j.biopsych.2012.06.006

Wong S, Le GH, Phan L, et al (2024) Effects of anhedonia on health-related quality of life and functional outcomes in major depressive disorder: A systematic review and meta-analysis. J Affect Disord 356:684–698. 10.1016/j.jad.2024.04.086

Xia Y, Driscoll JR, Wilbrecht L, et al (2011) Nucleus accumbens medium spiny neurons target non-dopaminergic neurons in the ventral tegmental area. J Neurosci 31:7811–7816. 10.1523/JNEUROSCI.1504-11.2011

Yalcin I, Belzung C, Surget A (2008) Mouse strain differences in the unpredictable chronic mild stress: a four-antidepressant survey. Behav Brain Res 193:140–143. 10.1016/j.bbr.2008.04.021

Yang H (2018) Nucleus accumbens subnuclei regulate motivated behavior via direct inhibition and disinhibition of VTA dopamine subpopulations. Neuron 97:434–449

Zhang C, Zhang C, Hong X, et al (2026) Cross−national comparison of major depressive disorder burden in China, India, and the United States of America: an age−period−cohort analysis of GBD 2021. Front Psychiatry 16:1686919. 10.3389/fpsyt.2025.1686919

Zhou K, Xu H, Lu S, et al (2022) Reward and aversion processing by input-defined parallel nucleus accumbens circuits in mice. Nat Commun 13:6244. 10.1038/s41467-022-33843-3

